# Convergent molecular evolution of viviparity across squamate reptiles

**DOI:** 10.1101/2025.09.26.678543

**Authors:** Hongxin Xie, Darren Hunter, John L Smout, Oscar E. Gaggiotti, Kathryn R Elmer

## Abstract

The molecular mechanisms underlying convergent phenotypic innovations across the tree of life are largely unknown. Viviparity (live-bearing) is a complex reproductive mode transition having more than 100 independent origins in squamate reptiles, making it an ideal model to understand the molecular basis of evolutionary innovations. Here we used 141 squamate genomes, including 42 viviparous species and representing more than 18 independent evolutionary transitions to viviparity, to search for convergent molecular signals in 14,096 protein coding genes. We show that convergent positive selection associated with viviparity at the gene-level and amino acid substitutions at the site-level are widespread in squamates, but there is no universal convergence across all viviparous clades. The chance of molecular convergence associated with viviparity is negatively correlated with genetic distance between species. Nevertheless, convergent genes are shared across independent origins of viviparity even spanning deep divergence times, and these genes are enriched in viviparity-related biological processes, including placental development, nutrient transport, and immune response to external organisms. Our result demonstrates that molecular convergence in the evolution of complex traits decreases in likelihood with evolutionary time, but when it occurs is more likely through changes in different genes of similar functional pathway rather than changes in the same genes.

## Main Text

A fundamental question in evolutionary biology is if phenotypic convergence is achieved by molecular convergence (1–3). Convergent phenotypes are a powerful insight into the shared and predictable nature of response to selection that occurs independently in different lineages. The extent of molecular convergence provides insight to the strength of selection and the flexibility of evolution to generate similar phenotypes from variable genetic backgrounds (4–7). The pattern of molecular convergence underlying a complex trait can be divided to at least three different levels: (i) homologous mutations at the same sites in the same genes across different taxa; (ii) different mutations in the same genes; (iii) different mutations in different genes that function in the same functional pathway and thus result in similar phenotypes (2, 7). The first two levels are generally considered rare, especially across highly divergent lineages, because of the low probability of targeted mutations at the same site and due to divergent genetic backgrounds on which mutations occur (i.e. epistatic constraint) (8). Epistasis is a pervasive phenomenon, in which the effect of a mutation is dependent on the interactions within the gene structure and between other genes. Consequently, the same novel mutation may not have the same functional outcome at higher biological levels in divergent lineages (1). Emerging genome resources have recently facilitated large scale genomic scans for convergent signals of trait evolution and have revealed cases of convergent gene mutations associated with ecologically relevant phenotypes (9–12), although other studies have failed to find significant molecular convergence (13–15). While phenotypic convergence is widespread across diverse taxa in nature, the extent to which it is underpinned by molecular convergence remains poorly understood.

Viviparity, or live-bearing, is a major reproductive innovation that has evolved independently across vertebrates, leading to new life history strategies and driving species diversification (16–18). While mammals are the animal group most well-known for giving birth to live young, in fact reptiles have most frequently transitioned from egg-laying (oviparity) to live-bearing. It has been estimated that there are more than 100 independent transitions to viviparity in squamates (lizards and snakes), compared to only once in mammals (19). Viviparity is a complex trait involving the modification of multiple functions to enable extended gestation time and development of the embryo in the maternal reproductive tract: enhanced nutrient, waste, gas, and water exchange between the mother and the embryo (often through the reduction of eggshell membrane and development of placenta), and consequently the suppression of maternal-foetal immune response as the developing embryo imposes extended exposure of foreign tissue within the mother. Finally, endocrine alterations in the maternal system are needed for longer embryonic retention (20–22). Thus viviparity is composed of a suite of biological changes working in tandem, with the shared outcome being the birth of free-living young.

Interest in dissecting the genetic and molecular mechanisms of viviparity has rapidly grown as the genomic resources for non-model species are emerging at pace. By comparing the gene expression in maternal reproductive tissue between pregnant and non-pregnant stages, and between oviparous and viviparous species pairs, studies have revealed that the expression of genes involved in viviparity can be shared across fish, squamates, and mammals, and many of the overlapping genes are involved in pathways related to tissue remodelling, nutrient transport, angiogenesis, and immune function (23–28). Nevertheless, efforts to find convergent changes in coding sequences (CDS) have been in vain, both in squamates (24) and fish (29). However, those studies were restricted to few transitions and in limited taxa, so low statistical power may have limited their findings. A more recent study spanning broad evolutionary time included 17 independent transitions to viviparity in fish (10 transitions), mammals (1 transition), and squamates (6 transitions) (30). Convergent changes in protein coding gene family sizes and CDS regions related to viviparity were identified in a handful of genes, but no convergent changes common to all viviparous lineages were found.

Here, we used broad scale genome sampling across the phylogenetic tree of squamates to include multiple independent transitions to viviparity with relatively consistent biological background in our dataset. We compiled, curated, and annotated a whole genome dataset for all squamates available to date: 141 genomes spanning 37 taxonomic families and including at least 18 independent origins of viviparity. We systematically scanned 14,096 orthologs for molecular signals of (i) positive - selection, (ii) intensified or relaxed selection, and (iii) convergent amino acid substitutions associated with viviparity. By comparing patterns across many independent viviparous clades, we tested how often and by what molecular routes a complex trait can evolve under divergent genomic contexts.

## Results

### Comparative genome resources and phylogeny

We collected 181 genomes for 161 different squamate species, as well as three outgroup reptile species from Rhynchocephalia, Testudines, and Crocodilia, all of which are oviparous (**table S1**). After filtering for genome quality and trait information (i.e. reported reproductive mode: oviparous or viviparous), a final list of 141 squamate genomes (140 different species and a subspecies of *Zootoca vivipara*) and the three outgroups were obtained (**table S2**). Our squamate genome dataset consisted of 37 families (representing 54.4% of all squamate families) and included 42 viviparous species from across 15 families. All genomes were annotated for protein-coding genes using a homology-based method, TOGA (31). The oviparous leopard gecko (*Eublepharis macularius*) genome and annotation were used as reference (32). Annotated one-to-one genes for each genome ranged from 5,804 to 16,528 (**Fig. 1, table S2**). To construct a species tree, 1,605 universal one-to-one genes for all species were extracted. The resulting phylogeny has high support for all major squamate clades (**Fig. S1**): Gekkota (geckos), Scincoidea (skinks), Lacertoidea (lacertids, spectacled lizards, whiptails and tegus, and legless worm lizards), Anguimorpha (anguimorph lizards such as monitor lizards, glass lizards, and alligator lizards), Iguania (iguanian lizards such as chameleons, agamas, iguanas, and anoles), and Serpentes (snakes). The main topologies we inferred are consistent with the recently published squamate phylogeny that was obtained with broader taxonomic coverage but fewer genetic markers (33). We used this topology as the species tree for all subsequent analyses.

**Figure 1.**
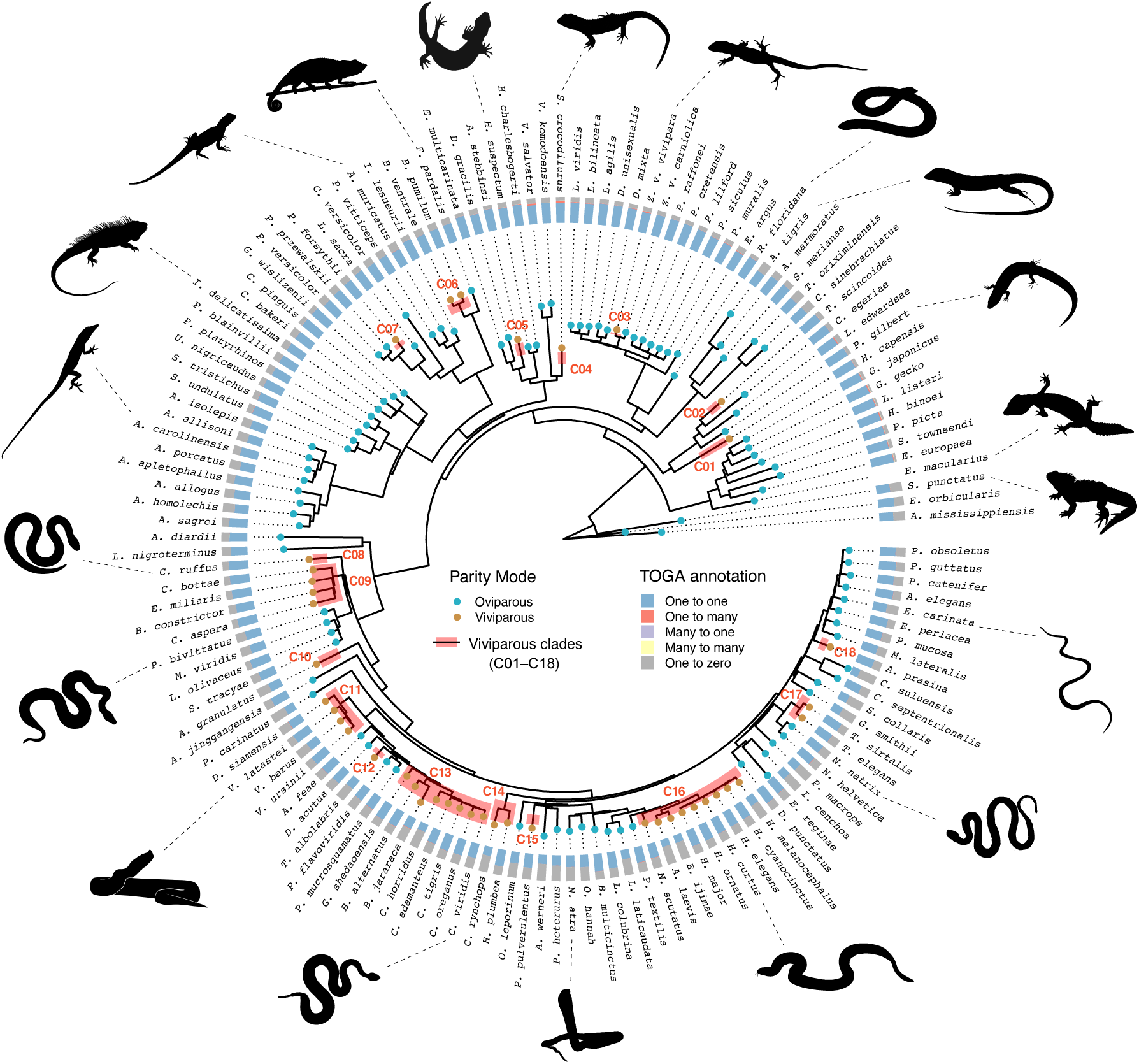
Species tree and genome annotation result. The squamate phylogenetic tree was built using maximum likelihood method based on protein alignment of 1,605 universal one-to-one genes. All major squamate clades received 100% bootstrap support. See **Fig. S1** for the detailed tree with bootstrap values. Eighteen viviparous clades were identified according to the phylogeny. Genome and protein-coding gene annotations of the leopard gecko (*E. macularius*) was used as the reference for gene annotation of all other genomes. Summary of TOGA annotation result for 20,456 genes are shown in a stacked bar plot. Animal silhouettes were collected from the open-access PhyloPic repository (https://www.phylopic.org).

Following established phylogenetic histories of reproductive mode evolution in squamates (18, 34, 35) and accepting the hypothesis that the ancestral reproductive state of squamate reptiles is oviparity (19, 36, 37), we resolved 18 different monophyletic viviparous clades in our species tree (C01–C18, **Fig.1**). Due to complex transition histories and lack of genome data availability for some taxa, not all oviparity-to-viviparity transition events can be captured in our phylogeny. For example, the clade comprised of pit vipers is completely viviparous in our dataset (C13), but oviparous species exist within this clade (38) that are not available to be included, meaning multiple independent transitions are collapsed as one transition. To account for such uncertainties, we focus our analysis on identifying convergent signals related to the evolution of viviparity across all viviparous branches in the phylogeny. Our analysis is not considering the reconstructed ancestral node where the oviparity-to-viviparity transition happened. Thus, our approach is conservative, in that we can only accurately estimate or underestimate (but not overestimate) independent transitions. Tolerating 50% missing data, a dataset of 25,464 transcripts from 14,096 genes was obtained for comprehensive searching of convergent signals in CDS regions. The dataset contains at least six independent viviparous clades for any gene alignment.

Having established a robust species tree and identified 18 independent viviparous clades, we used a phylogenetic framework of molecular evolution to test whether these lineages share signals of selection at the gene and site levels (**Fig. 2**).

**Figure 2.**
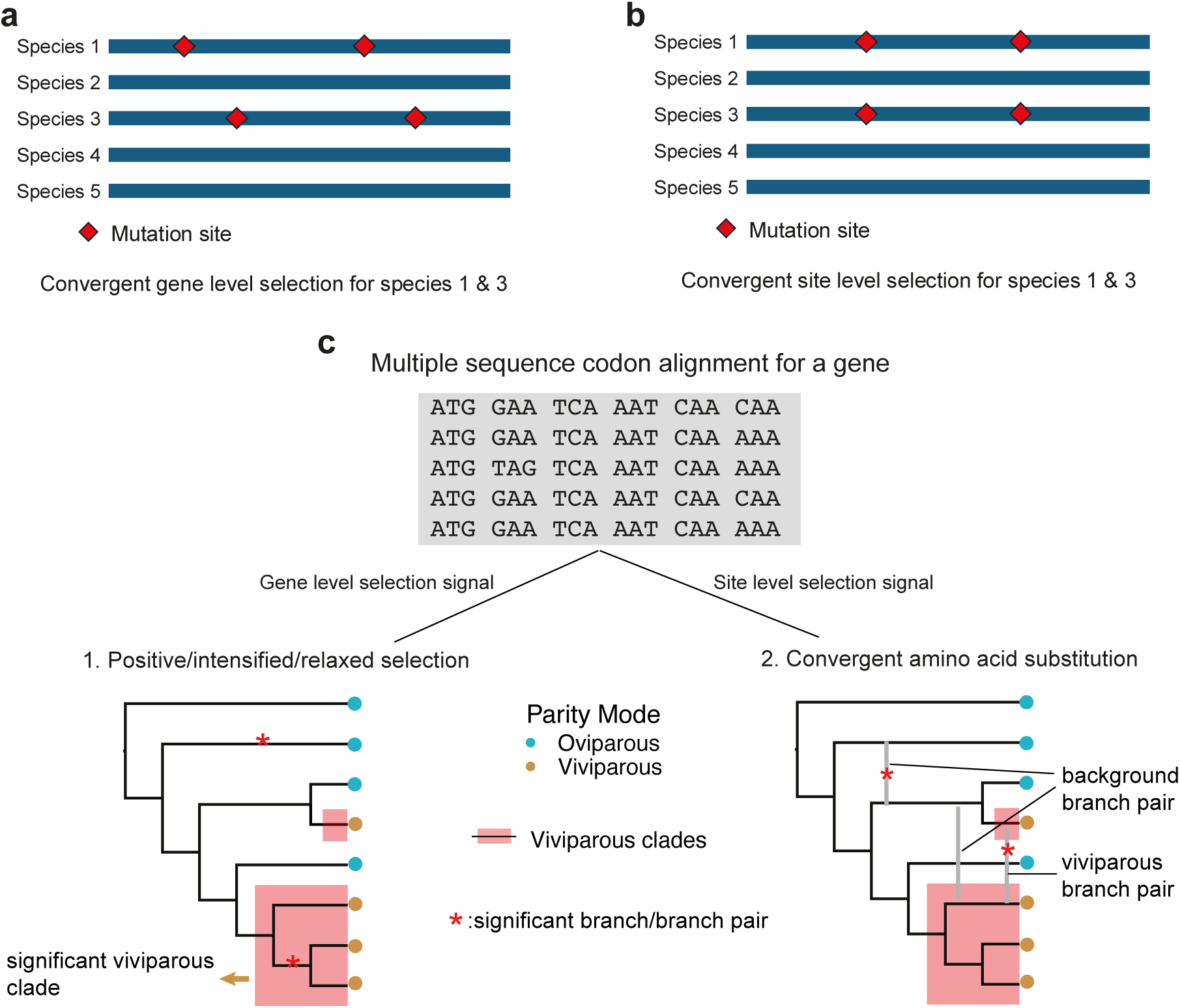
Schematic view of the analysis for detecting convergent selection signal for viviparity. (a) Convergent gene level selection signal. (b) Convergent site level selection signal.(c) The analysis conducted for each gene alignment. Analysis for gene level selection signal was conducted for each branch on the tree. A viviparous clade was considered significant if at least one internal branch showed significant signal (see Methods). For convergent substitution analysis, each branch pair was tested for convergence. Only viviparous-viviparous pairs were considered as foreground (viviparous branch pair). All other branch pairs (viviparous-oviparous and oviparous-oviparous) were considered as background. Branch pairs within the same viviparous clade were excluded.

### Convergent genes under selection in viviparity

The proportion of non-synonymous to synonymous nucleotide substitutions within a gene, relative to other features of evolutionary rate across a phylogeny, is a classical test of positive selection (39). An elevated non-synonymous to synonymous substitution ratio (*ω* = *d*_N_/*d*_S_; when *ω*>1) on specific branches indicates episodic diversifying selection (40). Here we focus on shared selection signals across the branches of multiple clades as evidence of convergent evolution in CDS regions, and we applied a series of codon-based methods (Fig. 2).

We first applied the aBSREL algorithm in HyPhy to detect positive selection across branches in the phylogeny (41). For each gene alignment in our dataset, each branch in the tree was tested and significant positively selected branches (PSBs) were obtained after correction for multiple testing. Out of the full dataset of 14,096 genes examined, no gene showed signals of positive selection across all 18 viviparous clades. Viviparous clades having at least one significant PSB for a gene ranged from 1 to 15, with 2,782 clade-specific positively selected genes (PSGs) but the gene count sharply decreased thereafter (Spearman’s rank correlation, *ρ* = –1, *p* < 0.0001; **Fig. 3a**). PSBs for 861 genes are found in viviparous branches but without signals of selection in the background oviparous branches; these are genes distinctly under selection only in viviparous clades. However, among these the maximum number of viviparous clades with PSBs (significant viviparous clades under positive selection) is four (**Fig. 3b**). Most genes have signals of positive selection both in the viviparous branches and in the oviparous background branches. These genes may be under positive selection for traits other than viviparity or relate to lineage-specific adaptations.

**Figure 3.**
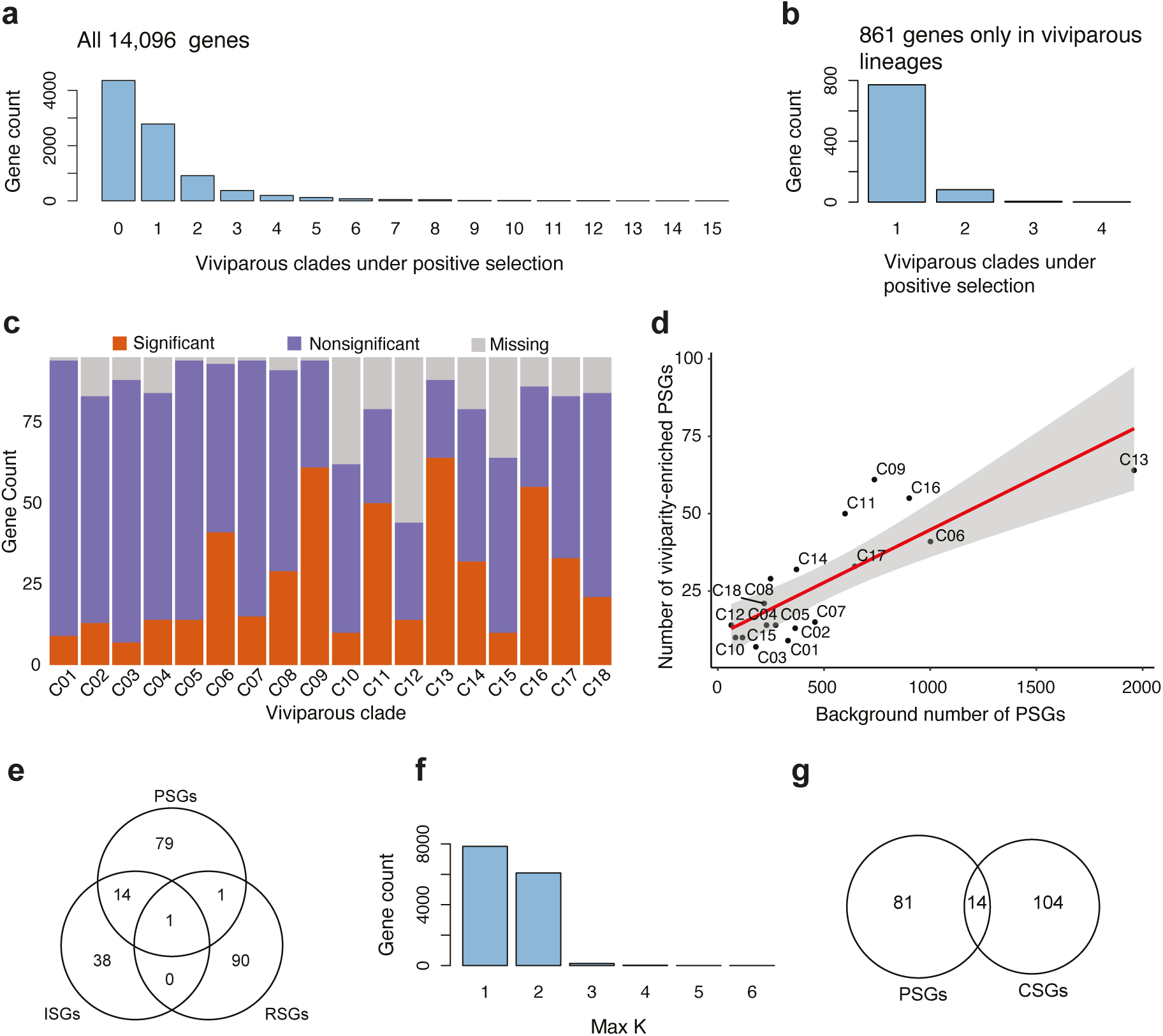
Results of selection signals in protein coding genes related to the evolution of viviparity. (a) Distribution of the number of significant viviparous clades under positive selection for all genes. For each gene, a significant viviparous clade is a viviparous clade with at least one internal branch showing signal of positive selection. (b) Distribution of the number of significant viviparous clades under positive selection for the 861 genes that exclusively show positive selection in viviparous lineages and not in oviparous lineages. (c) The number of genes showing positive selection signal in each viviparous clade for the 95 viviparity-enriched positive selected genes (PSGs). For each clade, the number of genes having “significant” positive selection signal, “nonsignificant” positive selection signal, or is “missing” in the alignment (due to assembly gap or quality control) are shown in stacked bar. (d) The correlation of the number of significant genes in the 95 viviparity-enriched PSGs and the number of background significant PSGs in each viviparous clade. The shaded area represents 95% confidence intervals of the regression line. (e) Venn plot showing the overlapping genes between the three viviparity-enriched gene sets: 95 positively selected genes (PSGs), 53 intensified selection genes (ISGs) and 92 relaxed selection genes (RSGs). (f) Distribution of the maximum number of convergent clades (max *K*) detected for amino acid substitutions for all genes. Max *K* =1 indicates no convergence in amino acid sites found across all branch pairs for the gene. (g) Venn plot showing the overlapping genes between the 95 viviparity-enriched positively selected genes (PSGs) and 118 viviparity-enriched convergent substitution genes (CSGs).

To focus on genes under selection that are specifically related to viviparity, we defined a “viviparity enrichment factor” for genes that have a higher proportion of PSBs in viviparous branches relative to the background (42). We identified 232 positively selected genes (PSGs) that show enriched signal in viviparous branches (enrichment factor >1, contingency table *p* < 0.05, Fisher’s exact test; **table S3**). After false discovery rate (FDR) control, 95 genes were identified as significant viviparity-enriched PSGs (**Fig. S2**, **table S3**). This set of 95 genes is widely shared and found across all 18 viviparous clades (**Fig. 3c**). Nonetheless the convergence at gene level is not extensive: 78.9% of PSGs (genes with PSBs) are found across six or fewer viviparous clades and no single gene is found in more than 15 clades (**Fig. S3**). The number of viviparity-enriched PSGs in each viviparous clade is positively related to the background number of PSGs (**Fig. 3d, Fig. S4**) and there are no extreme outliers for the regression model (largest studentized residual = 2.65, Bonferroni-adjusted *p* =0.324), suggesting the convergent signal that is present is not dominated by specific clades.

Selection pressures on genes related to a trait innovation can become stronger (intensified selection) or weaker (relaxed selection) in lineages of interest (43, 44). We expect viviparity-enriched PSGs are also under intensified positive selection in viviparous lineages, but not under relaxed selection. To detect intensified or relaxed selection in the phylogeny, we applied RELAX in HyPhy (45) and - similar to our approach for inferring positive selection - each branch was tested and enrichment inferred relative to oviparous background. We identified 205 genes showing signals of intensified selection (ISGs: intensified selection genes) that were enriched in viviparous branches (enrichment factor >1, contingency table *p* < 0.05, Fisher’s exact test; **table S4**). Fifty-three viviparity-enriched ISGs were retained after FDR control (**table S4, Fig. S5**). 81.1% of viviparity-enriched ISGs were found to be significant across six or fewer viviparous clades (**Fig. S6**). This is a similar level of convergence as we had found with the viviparity-enriched PSGs. Fifteen genes were shared between the viviparity-enriched PSGs and viviparity-enriched ISGs, showing significantly more overlap than expected by chance (*p* < 0.0001, hypergeometric test; **Fig. 3e**). This supports the expectation that genes under positive selection for viviparity are experiencing intensified selection in viviparous lineages.

We found 212 genes showing signals of relaxed selection (RSGs: relaxed selection genes) that were enriched in viviparous branches (enrichment factor >1, contingency table *p* < 0.05, Fisher’s exact test; **table S5**) and 92 viviparity-enriched RSGs were retained after FDR control (**table S5, Fig. S7**). The set of viviparity-enriched RSGs has a higher convergence level than we found in ISGs, with 79.3% of genes found in up to 10 viviparous clades having PSBs (**Fig. S8**). In contrast to the significant overlap of positive and intensified selection (PSGs and ISGs), only two genes were shared between the viviparity-enriched PSGs and viviparity-enriched RSGs, which is not different from chance (*p* = 0.28, hypergeometric test; **Fig. 3e**). This demonstrates that relaxed selection affects a distinct set of genes and is not interplaying with positive selection in viviparity.

### Convergent site substitutions under selection in viviparity

These tests of convergent PSGs we conducted consider signals summarised across the entire CDS (**Fig. 2**), but it can be that selection is even more focused, targeting amino acid sites across independent viviparous clades and driving convergent amino acid substitutions in particular sites (46, 47). This is important and complementary because convergent substitutions at the same residues can reveal parallel adaptive solutions at functionally critical sites, which might be obscured when averaging signals across whole genes (12). To search for convergent amino acid substitutions at the same specific sites in a gene between branch pairs, we applied CSUBST, a method that incorporates combinatorial substitution ratios (*ω_C_*) to account for background genetic noise and phylogenetic errors (48). CSUBST detects convergent events in the foreground (here, viviparous clades) to the maximum number of convergent clades (*K*) in which they exist. Using a cut-off of *ω_C_* > 3 and requiring at least two convergent amino acid sites (which is less stringent than the default setting but still accounts for random homoplasy), the maximum convergence obtained for any gene tested in our dataset was six (i.e. maximum *K* = 6) (**Fig. 3f**). The vast majority of genes (97.5%; 5,887 of 6035 genes) showing a convergent signal at the same amino acid sites were found only between two clades (i.e. have maximum *K* = 2). This demonstrates that convergent amino acid substitutions across independent viviparous clades are extremely rare.

Focusing on convergent signals found across two clades (*K* = 2), we again defined an “viviparity enrichment factor” to find genes that have higher proportion of branch pairs showing convergent substitution signals in viviparous clades compared to background. We obtained 211 convergent substitution genes (CSGs) that show enriched signal in viviparity (enrichment factor >1, contingency table *p* < 0.05, Fisher’s exact test; **table S6**). Controlling for false positives, we accepted 118 viviparity-enriched CSGs (**table S6**). Of this gene set, 14 are also enriched for positive selection in viviparous lineages in our previous analyses, showing a significant overlap (*p* < 0.0001, hypergeometric test, **Fig. 3g**). This demonstrates that a significant portion of the viviparity-enriched PSGs also show (or are specifically due to) convergent amino acid substitutions that are driven by positive selection.

### Molecular convergence is negatively related to phylogenetic distance

We have detected widespread convergent selection signals among sets of viviparous clades, although with a lack of high-order convergence shared across most or all clades of squamates. If epistasis explains our observed pattern, we expect to see convergence between two independent clades decrease with their phylogenetic distance increasing (1). Consistent with this hypothesis, we observed a negative correlation between the number of viviparity-enriched PSGs shared between two viviparous clades (scaled using Jaccard similarity) and the pairwise phylogenetic distance between those clades (*r* = -0.44, p < 0.0001; **Fig. 4a**). The background gene set (i.e. convergent genes not enriched for viviparity) also shows a negative correlation between shared PSGs and pairwise phylogenetic distance, but to a lesser extent (*r* = -0.22, p = 0.0060; **Fig. 4b**), demonstrating that phylogenetic distance imposes a stronger negative effect on the sharing of PSGs in viviparous clades than on background convergence across squamates.

**Figure 4.**
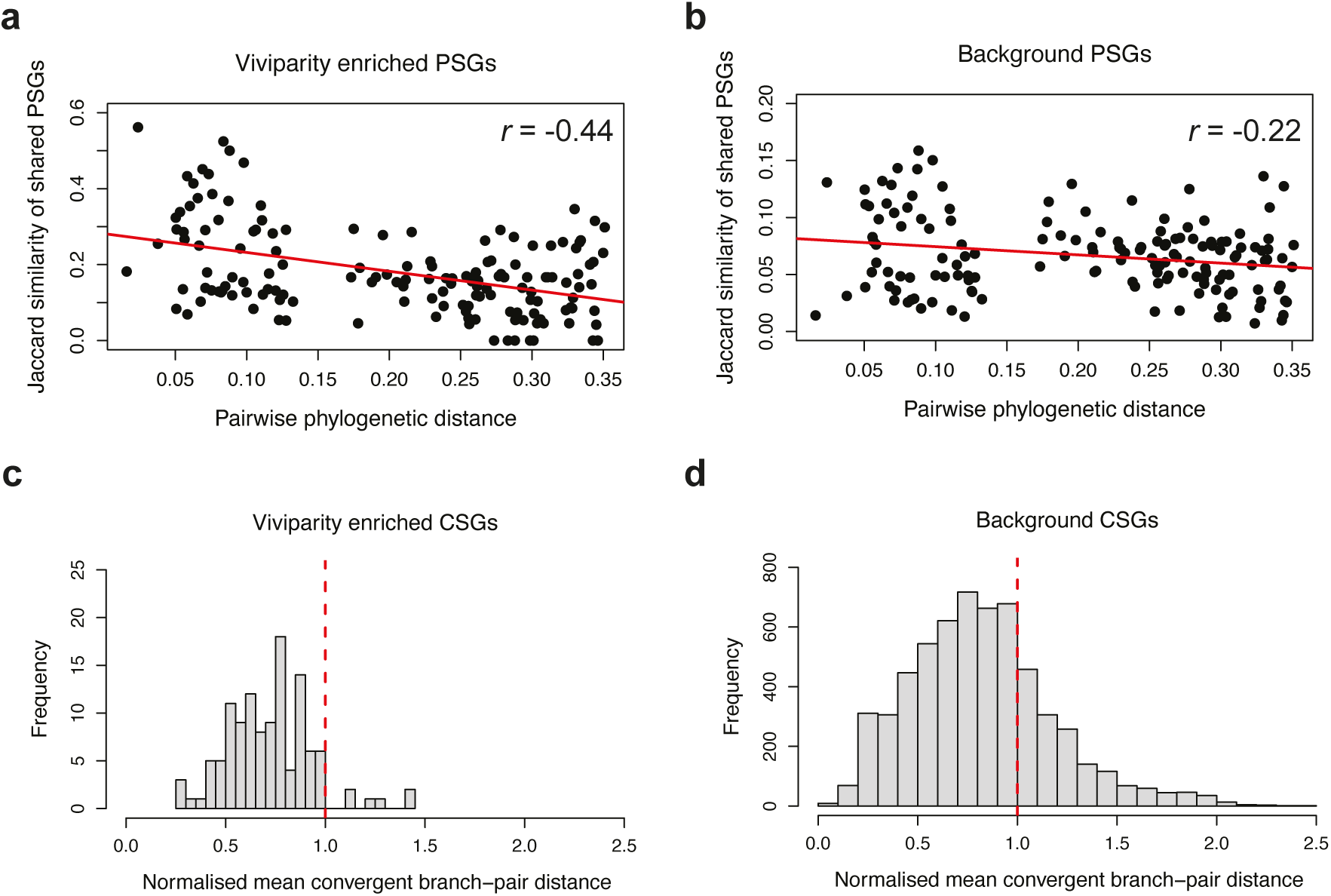
The relationship between the frequency of molecular convergence and phylogenetic distance. (a) & (b) The regression of pairwise distance of shared number of PSGs (quantified using Jaccard similarity) and their phylogenetic distance. Plots were drawn for the 95 viviparity-enriched positively selected genes (PSGs) and all the other background PSGs, respectively. (c) & (d) Distribution of the mean paired-branch distance for the viviparous lineages showing convergent substitution signal. Convergent distance was normalised by the distance of branch pairs without convergent signal. The red dashed vertical line represents the null hypothesis of a normalised distance of “1”. Plots were drawn for the 118 viviparity-enriched convergent substitution genes (CSGs) and all the other background CSGs, respectively.

For amino acid substitutions, we identified CSGs by finding convergent events between branch pairs in the viviparous foreground. To test if the convergent events are influenced by the evolutionary distance of the paired branches, for each gene we calculated the mean distance between the branch pairs that show convergent signal in viviparous foreground. We normalised the distance between pairs of convergent branches by the distance of nonsignificant branch pairs in the viviparous foreground. The null hypothesis would be a mean distance of 1, where phylogenetic distance has no relationship with site convergence. Alternatives include <1-shifted, where convergence is more likely between closely related clades, or >1-shifted, where convergence is more likely between distant clades. The 118 viviparity-enriched CSGs show a distribution of normalised mean convergent branch-pair distance biased towards 0 (median = 0.73, *p* < 0.0001, Wilcoxon signed-rank test; **Fig. 4c**). The background gene set (the genes that have convergent branch pairs but not enriched in viviparity) also show a distribution of normalised mean convergent branch-pair distance biased towards 0 but to a lesser extent (median = 0.79, *p* < 0.0001, Wilcoxon signed-rank test) and the mode is shifted closer to 1 (**Fig. 4d**). Only 5.1% of viviparity-enriched CSGs have a normalised mean convergent branch-pair distance >1, while 26.2% of background convergent genes have a normalised distance >1. This demonstrates a stronger negative effect of phylogenetic distance on site convergence in viviparous clades compared to background convergence events.

Combined our results demonstrate that phylogenetic distance negatively influences the chance of convergence under positive selection for independent viviparous clades, both at the gene level and at the level of amino acid substitutions.

### Functional enrichment of convergence

We predicted that the set of genes we have identified as responding in a convergent manner to positive selection in viviparous clades are highly likely to have functions associated with live-bearing traits. To test this and identify the biological processes these convergent genes are involved in, we performed Gene Ontology (GO) analysis and curated candidate pathways for viviparity. We found the 95 viviparity-enriched PSGs are enriched in 25 GO terms (**Fig. 5a, table S7**), including biological processes related to placenta development (GO:0060710: chorio-allantoic fusion, GO:0048608: reproductive structure development), nutrient transport (GO:0044539: long-chain fatty acid import into cell, GO:0006904: vesicle docking involved in exocytosis), and immune response to external organisms (GO:0071219: cellular response to molecule of bacterial origin). These GO terms are notably consequential for reproduction, pregnancy, and foetal-membrane development (**Fig. 5b**).

**Figure 5.**
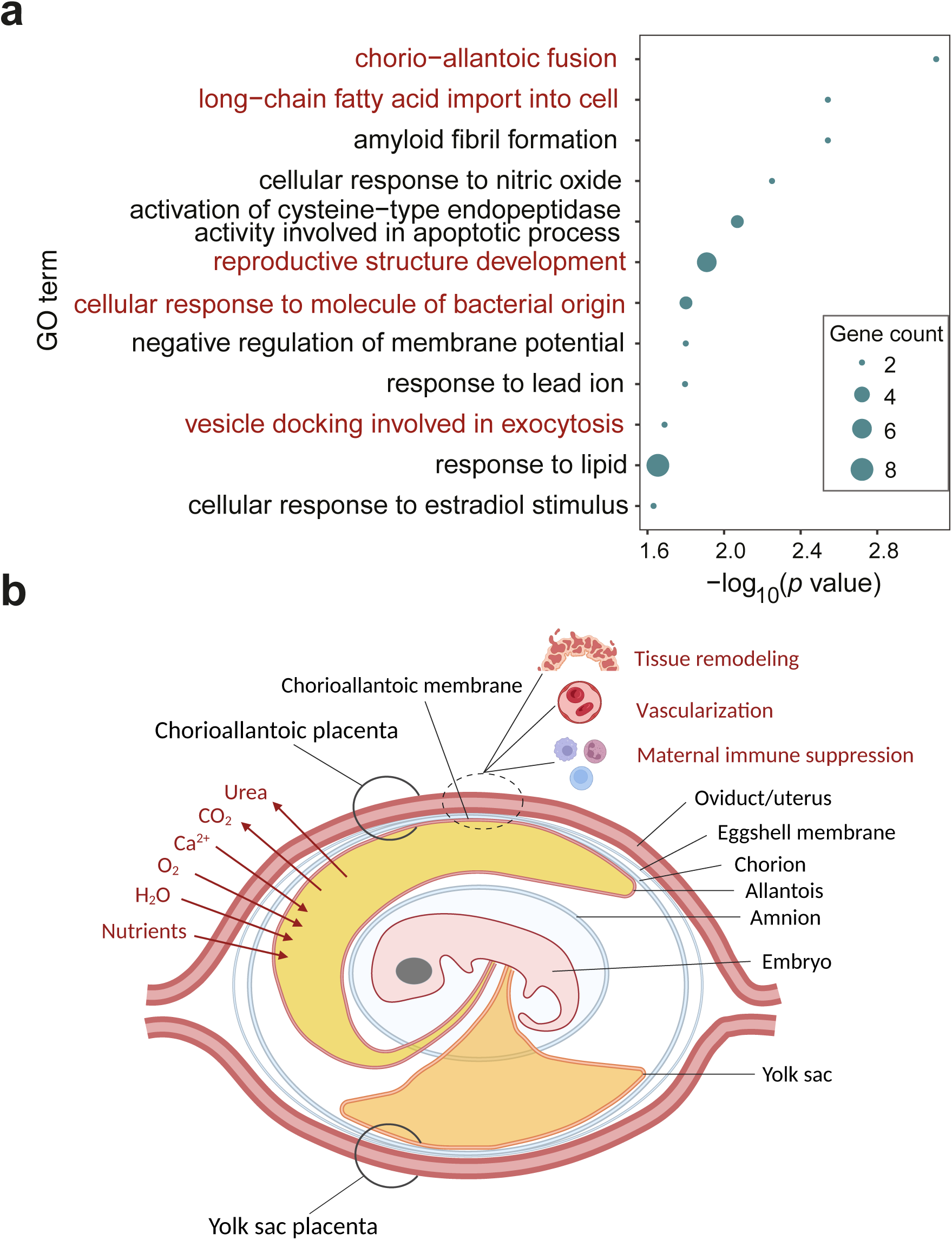
Enriched functional pathways of viviparity-enriched genes. (a) The top 12 significant Gene Ontology terms for the 95 viviparity-enriched positively selected genes (PSGs). See **table S7** for the full list. Terms having a direct relationship with viviparity are in red. (b) Schematic illustration of the major physiological functions needed for live-bearing. Chorioallantoic placenta is shown as an example, but note that viviparous squamates generally develop both a chorioallantoic and a yolk-sac placenta to various degrees (26). Structures are in black font and functions are in red. Illustration was drawn in BioRender

Individual viviparity-enriched PSGs we identified have known functions in the development of mammalian placenta. For example, *ERF* is transcriptional repressor required for extraembryonic ectoderm differentiation, ectoplacental cone cavity closure, and chorio-allantoic attachment (49). *ITGA4* encodes the integrin subunit alpha 4, a crucial cell surface protein involved in cell adhesion and signaling, and has been shown to function in mouse, ovine, and human placenta development (50–52). *BPTF* expression is essential for trophoblast differentiation in early pregnancy (53, 54).

Similar to the pattern inferred from PSGs, the 53 viviparity-enriched ISGs are also significantly enriched in terms related to placenta development (GO:0001893: maternal placenta development), nutrient homeostasis (GO:0070328: triglyceride homeostasis, GO:0042593: glucose homeostasis), and immune responses (GO:0071219: cellular response to molecule of bacterial origin, GO:0042119: neutrophil activation; **table S8**). In contrast, enriched biological processes for the 92 viviparity-enriched RSGs show little relationship with live-bearing (**table S9**), a pattern which further bolsters the biological relevance and resolution of the convergent genes under positive selection.

The 118 viviparity-enriched CSGs we found are enriched in 45 GO terms (**table S10**). The enriched biological processes are dominated by immune responses, and also include functions related to nutrient transport (GO:0034197: triglyceride transport, GO:0008645: hexose transmembrane transport, GO:0051223: regulation of protein transport) and regulation of metabolism (**table S10**), all of which are relevant to live-bearing (20–22).

We have also identified 14 genes that belong both to viviparity-enriched PSGs and viviparity-enriched CSGs (**Fig. 3g**). These genes not only show shared positive selection signal at gene level across independent viviparous clades (**table S11**) but also contain convergent amino acid site substitutions; these represent a short-list of strong candidates with molecular adaptations in response to the evolution of viviparity. Among them we found the genes *IL6* and *CD36* to be particularly involved in multiple functional terms in the enriched function list for viviparity enriched PSGs (**table S7**) and CSGs (**table S10**). *IL6* is a cytokine (interleukin-6) with broad function in immunity, inflammation, tissue regeneration, and metabolism (55). Normal expression of *IL6* in human placenta promotes angiogenesis and remodelling of endometrial blood flow, and maintains immune homeostasis during pregnancy (56–58). *CD36* is a multifunctional glycoprotein (platelet glycoprotein 4) that couples lipid metabolism to inflammatory, immune, and angiogenic signalling across many tissues (59). In the mammalian placenta, *CD36* is polarized to syncytiotrophoblast membranes where it governs fatty-acid transport and plays an essential role for successful pregnancy (60, 61). We consider these two genes to be very strong candidates for molecular adaptation linked to evolution of viviparity.

Because enrichment analyses based on predefined gene sets depend on a significance threshold, they may overlook pathways composed of many genes with moderate but coordinated signals. To complement this approach, we applied Gene Set Enrichment Analysis (GSEA), which evaluates all genes ranked by selection signal and provides a threshold-free perspective on biological pathways under selection for viviparity. A total of 333 KEGG pathways were mapped to the 14,096 genes used in selection analyses. Top pathways enriched for positive selection and intensified selection are related to energy metabolism (ko00071: fatty acid degradation, ko00190: oxidative phosphorylation), immune response to pathogens (ko04666: Fc gamma R-mediated phagocytosis, ko04621: NOD-like receptor signalling pathway), and many modules associated with cancer biology (**table S12&S13**). While pathways under relaxed selection are dominant in basic physiological processes different from those observed in positive and intensified selection (**table S14**). Top pathways under convergent amino acid substitutions are also dominated by infectious disease and oncogenic signalling modules (**table S15**). Cancer-related pathways enriched in our analyses likely reflect the activation of core cellular programs such as proliferation, invasion, angiogenesis, and immune modulation, which are not unique to tumors but are also central to placental development and the extensive tissue remodelling required for viviparity (62–64).

## Discussion

Our evolutionary analysis of 141 squamate genomes, capturing 18 independent clades of live bearing across lizards and snakes, shows that viviparity repeatedly recruits the same physiological pathways (placental formation, nutrient transport, and immune modulation) but almost never the same exact genetic changes. Selection is therefore highly predictable at the level of biological process, yet strikingly varied at the level of genes and amino-acid sites. This dual pattern resolves the long-standing paradox of widespread phenotypic convergence in the evolution of vertebrate reproduction alongside very little evidence for universal molecular convergence (24, 29, 30)

Our study leverages the largest comparative genomic dataset to date for reptiles. Drawing on the recent growth in publicly available genomic data, we have dramatically increased sample size compared to previous efforts (24, 29, 30). This expanded sampling not only increases statistical power to detect subtle signals of convergence, but also provides a phylogenetic framework to robustly disentangle lineage-specific adaptations from shared evolutionary pressures. By applying a uniform reference-based gene annotation, we obtained 14,096 one-to-one gene alignments, which is also by far the largest to date for squamates (24, 30). Codon alignment-based selection scans are known to be affected by the quality of the alignment, with poor alignment leading to high false positive rate (65, 66). We made use of a state-of-the-art homology-based method to accurately separate orthologues from paralogues and conduct scaled gene structure annotation (67). The large-scale alignments obtained here allowed us to conduct an accurate and comprehensive scan for convergent selection signals associated with viviparity.

Signals of positive selection, intensified selection, and convergent amino acid substitutions are widespread among viviparous lineages, yet rarely shared by more than a few clades. Across the genes enriched for positive selection and/or intensified selection in viviparous lineages, no single locus shows signal of selection in all 18 viviparous clades, and the vast majority of the genes appear in only a few clades (**Fig. 3b, Fig S6**). Similarly, of those genes with enriched convergent amino acid substitutions in viviparous lineages, almost all convergent substitutions occur in just one pair of viviparous clades (i.e. *K* = 2). These results reinforce the view that while evolution can ‘re-use’ the same genes under selection, it rarely does so in exactly the same way across deeply diverged clades (2, 68). Instead, convergent trajectories appear largely pairwise and patchy, consistent with an epistatic model that the core “viviparity toolkit” is deployed selectively in a lineage-specific manner rather than universally. Similarly, previous findings on the convergent regulatory change signals for viviparity also revealed no universal differential expression of any gene across viviparous amniotes (69), although regulatory convergence within a more limited number of taxa has been reported (23). Our results on this comprehensive set of protein-coding genes further confirms the absence of a molecular change linked with viviparity that spans reptiles.

The negative relationship between phylogenetic distance and convergent molecular signal both in positive selection (**Fig. 4a**) and amino acid substitutions (**Fig. 4c**) is robustly demonstrated in our dataset. We conclude that this indicates that deeper divergence imposes stronger epistatic constraints on repeatability. In other words, the more distantly related two viviparous lineages are, the less likely they are to arrive at the same genetic changes. This pattern held even among genes not associated with viviparity, albeit with weaker effect sizes (**Fig. 4b & 4d**). Phylogenetic constraints on neutral molecular convergence rate have long been proposed based on mitochondrial and nuclear genomic data (1, 70–72). Beyond confirming the general principle of divergence-dependent constraint on molecular convergence, our results shows that under adaptive selection for viviparity, epistasis effect is not weakened but instead reinforced for the recruitment of viviparity-related gene mutations. We speculate that this increase in epistasis effect and decreased molecular convergence with increasing phylogenetic distance might be generally applicable to other complex, polygenic traits. This remains to be evaluated in future studies.

Despite there being limited overlap at the level of individual genes or amino acid sites, our functional enrichment analyses reveal coherent molecular convergence at biological processes related to live-bearing (**Fig. 5b**). Positively selected and intensified selection genes are enriched for nutrient transport, immune response to external organisms, and placenta-related developmental pathways. These are all functions critical for extended gestation and nutrient exchange (20–22). Convergent substitution genes mainly cluster in immune related pathways, pointing to repeated evolution of maternal-foetal immune modulation to facilitate extended gestation (20). These enriched biological processes for viviparity are also frequently enriched in differential expression studies on the reproductive tissue of viviparous females (23–26, 69). GSEA analysis further detected common pathways related to oncogenic processes that are under convergent positive selection for viviparity, pointing to a well-recognized theme that tissue remodelling process in placentation co-opts pathways associated with tumor biology (62–64). Thus, while the precise genetic modifications differ, viviparous squamates appear to target the same functional modules to solve common physiological challenges in the evolution of live bearing.

Our findings suggest a hierarchical model of evolutionary predictability for a complex trait like viviparity: high predictability at the pathway level, where selection repeatedly recruits variation into a finite set of physiological functions shared for live birth (20); lower predictability at the gene or site level. Certain genes are more likely than random to be recruited, yet rarely across more than a handful of lineages. Individual amino-acid changes can recur, but seldom beyond a pair of branches. This result underscores that while natural selection can shape similar phenotypes through convergent pressures, the molecular routes are constrained by each lineage’s genomic background and historical mutations.

Research on convergent phenotypes - from fish ecotypes to flower color - have previously suggested a lack of molecular convergence at the genome level but increasing convergence at the level of pathways (73–75). What is notably different about viviparity from other complex traits is that it is widely regarded as a step-change in evolution that is lacking intermediate phases from oviparity (36, 76). Building on this framework, future work should explore regulatory and non-coding landscapes in the genome, which may be more pervasive but also more challenging to detect. The candidate genes we identified in this study (e.g., ERF, ITGA4, IL6) merit functional assay validation across squamate models to better understand amniote viviparity and placentation. Moreover, squamates exemplify a genomic “natural experiment” for studying repeated complex trait evolution. Similar comparative genomic approaches for other repeatedly evolved traits in squamates (e.g., limb reduction, arboreal adaptation) and convergent traits in other taxa with rich genome resources could reveal whether the hierarchical predictability we observe here is a general feature of evolutionary innovation. In summary, by combining an unprecedented squamate genome resource with codon-based selection scans and convergent-substitution analyses, we demonstrate that complex traits like viviparity evolve through a mosaic of shared functional pressures and lineage-specific molecular solutions. This comprehensive picture of convergence advances the understanding of evolutionary predictability and highlight squamates as a natural model for dissecting the molecular mechanisms of repeated phenotypic innovations.

## Methods and materials

### Data collection and curation

We exhaustively collected squamate genomes from all available public repositories, including published and unpublished ones (**Fig. S1**). Genomes were updated until February 2024. Species phenotypes (oviparous or viviparous) were collected from reported information in literature, here mainly from the attached phenotype information in two articles (18, 77). Collected genomes were then curated for quality. This study focuses on protein-coding genes; we therefore filtered genomes mainly based on the Benchmarking Universal Single-Copy Orthologs (BUSCO) scores (78). BUSCO scores were obtained for each genome assembly using BUSCO v5.2.2 and the sauropsida_odb10 database (7480 genes). Genomes with < 50% BUSCO for complete genes or > 10% BUSCO for complete duplicate genes were excluded. An extremely high duplication rate indicates possible assembly errors (79) or species-specific genome characteristics, both may cause bias to the dataset. Duplicated genomes for the same species were removed so that one genome with best BUSCO scores was retained. Species with no definite reproductive mode information were also removed.

### Genome annotation

Part of our collected genomes have protein-coding gene annotation files available in NCBI or from the authors, but a large number of genomes don’t have annotations. To obtain structural annotation for all genomes and avoid possible systematic bias if we incorporate annotations from multiple resources, we annotated every genome using TOGA v1.1.5, a recently developed gene-homology-based method that has been shown to be accurate and powerful (31). TOGA integrates structural gene annotation and orthology inference based on a single reference genome. We chose the leopard gecko, *E. macularius*, as the reference because it’s among the genomes with the best completeness in squamates, has an up-to-data annotation from NCBI, and is the ancestral phenotype (oviparous).

To facilitate annotation, genomes were masked for repeat content. Repeat sequences in the genome were first *de novo* identified by RepeatModeler v2.0.4 (80) using “-LTRStruct - quick” options to include LTR retrotransposons and use the reduced sample sizes of the quick mode because the full default pipeline takes extremely long for a large vertebrate genome. Genomes were then masked using the identified repeat families by RepeatMasker v4.1.5 (81) using “-norna -xsmall” options to skip masking non-coding RNAs and conduct soft masking. TOGA annotation was conducted in a two-step manner. Query genomes were first aligned to the reference genome using “make_chains.py” v1.0.0 (https://github.com/hillerlab/make_lastz_chains) to generate pairwise chain files. Structural annotation was then done by TOGA with gene-transcript information of the reference genome annotation provided (34,175 transcripts from 20,456 protein coding genes) and with “--ms” option to mask stop codons in target sequences.

### Species tree construction

We used strict universal one-to-one genes across all species for constructing the species tree. Orthology information was extracted directly from TOGA results. Extracted CDS were processed by the “codon-msa” pipeline from HyPhy-analyses repository (https://github.com/veg/hyphy-analyses/tree/master) to obtain in-frame codon alignments.

Briefly, extracted nucleotide sequences for each gene was first processed by “pre-msa.bf” to correct for frameshift mutations and generate protein sequences. Protein sequences were then aligned using MAFFT v7.505 (82) with default parameters. Protein alignments were back translated to codons using “post-msa.bf”. Finally, a custom python script was used to mask all invalid codons in the alignment to “---” (a valid codon should have “A”, “T”, “G”, or “C” in all three positions). The fixed in-frame codon alignments were further trimmed by trimAl v1.2rev59 (83) using “-automated1” option to remove poorly aligned columns. Gene alignments were concatenated using AMAS (84). The concatenated protein alignment was used to build a maximum likelihood tree by iqtree v1.6.12 (85) with 1000 ultrafast bootstraps and the “-spp” option for partitioned analysis. Based on the species tree, viviparous tips as well as all their internal nodes were considered viviparous lineages based on maximum parsimony principle and the assumption of no reversal.

### Exon-by-exon alignments of protein-coding genes

To gain as many one-to-one orthologue genes as possible for selection analyses, we extracted codon alignments for each transcript directly from TOGA annotations using the “extract_codon_alignment.py” script provided in TOGA repository (31). Only one-to-one orthologues were aligned (“--skip_dups”) and other parameters were set as default. By default, the script extracts orthologous sequences from a list of genomes according to the reference transcript sequence and conducts exon-by-exon alignment using MACSE v2.07 (86). Alignments were then filtered by HmmCleaner v0.243280 (87) to remove poorly aligned residues. To that end, each alignment was passed through a final fixing step by a custom python script to mask all invalid codons in the alignment to “---”. Finally, alignments with more than 50% missing data (72 sequences) were excluded for all downstream analysis. The accepted dataset includes 25,464 transcript codon alignments from 14,096 genes. All selection analyses were all based on these curated codon alignments.

### Positive selection analysis

We used a branch-site model, aBSREL in HyPhy v2.5.59, to scan for branches showing signals of positive selection (41). Our phylogeny includes multiple separate origins of viviparity and we aimed to detect genes that are under positive selection exclusively in viviparous lineages, or show enriched selection signal compared to oviparous background. We therefore ran aBSREL in explosive mode for each transcript alignment, testing every branch on the tree for positive selection. Only branches with an adjusted *p* value < 0.05 (Holm-Bonferroni correction) were accepted as showing significant signals of positive selection. We then defined a viviparity enrichment factor, which was inspired by the “relevance ratio” used in a previous study (42), as:

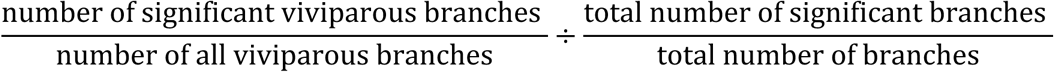

An enrichment factor > 1 indicates that a higher proportion of branches show positive selection in the viviparous lineages. A contingency table following Fisher’s exact test was applied for each transcript alignment to test if this difference in proportion is significant. Our dataset includes isoforms for genes. The isoform with the lowest *p* for the enrichment factor was chosen to represent each gene. Under this framework, genes with significant enrichment factor > 1 were considered viviparity-enriched PSGs. To control for FDR while retaining the most statistical power, we applied 1,000 empirical permutations of the foreground nodes on the tree. For each permutation, the same number of viviparous nodes were assigned randomly on the species tree. We then calculated the *p* value for each gene using the same Fisher’s exact test. For each original observed *p* value (*t*) between 0 and 0.2, the empirical FDR was obtained by the expected number of false positive genes obtained from permuted dataset (the mean number of genes having permuted *p* ≤ t) divided by the observed number of positive genes (number of genes having observed *p* ≤ t). We accepted an exploratory FDR threshold of 0.25 in this study to get a confident candidate list of convergent genes for downstream analyses. This threshold is commonly used for studies exploring functional pathways of candidate gene set (88, 89). However, this threshold is intended for exploratory assessment of functional relevance and should not be considered decisive for identifying individual genes. We also report the list of genes at FDR threshold 0.1 and 0.2 (**table S3**).

### Intensified and relaxed selection analysis

Complementary to aBSREL, another model in HyPhy, RELAX (45), was used to detect signals of intensified or relaxed selection in the phylogeny. RELAX utilises a selection intensity parameter (*k*). A significant *k* > 1 indicates that selection strength has been intensified on the tested branch, and a significant *k* < 1 indicates relaxed selection strength (45). We used a modified version of RELAX (https://github.com/veg/hyphy-analyses/tree/master/RELAX-scan) to scan for signals of intensification/relaxation for every branch individually on the tree relative to the gene background. Similar to aBSREL, only branches with an adjusted *p* value < 0.05 were accepted as having significant intensified/relaxed selection. We also calculated the viviparity enrichment factors for intensified selection and relaxed selection, respectively, in the viviparous lineages. Contingency table tests were applied to each transcript to test if the enrichment factor > 1 was significant. The isoform with the lowest *p* for the enrichment factor was chosen to represent each gene. Finally, genes with significant enrichment factor for intensification >1 were considered viviparity-enriched ISGs; genes with significant enrichment factor for relaxation >1 were considered viviparity-enriched RSGs. The same empirical permutation-based FDR analysis conducted for PSGs were also applied for ISGs and RSGs.

### Convergent amino-acid substitution analysis

We used a recently developed method, CSUBST (48), to detect adaptive convergent amino-acid substitutions for viviparity. CSUBST developed a new parameter, the combinatorial substitution ratios (*ω_C_*), to distinguish true adaptive convergence from random neutral convergent substitutions. This new error-corrected parameter has been shown to be powerful and accurate, and can tolerate phylogenetic errors (48). We ran CSUBST analysis by setting the viviparous clades as foreground lineages and set “--fg_stem_only no” to consider pairing branches for all internal nodes in the viviparous foreground but not only considering the stem node for each viviparous clade. Branch pairs within the same viviparous clade were excluded for the analysis. The analysis starts by finding convergent signal between two branches (*K* = 2). Higher order convergence was searched for based on the result of prior round (progressive heuristic algorithm). The program will progressively increase *K* until no significant convergence was detected. In theory, a branch pair of *ω_C_* > 1 and with more than one convergent nonsynonymous mutations 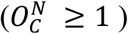 indicate adaptive convergent amino-acid substitutions, but a cut-off of 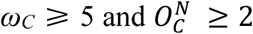 was recommended in the original paper because a single site convergence between only two lineages may also be from random homoplasy (48). In this study, we set the cut-off to 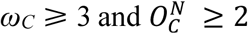 to both controlling for random error and trying to detect more subtle adaptive convergence signal across our multiple independent lineages of viviparity. We define a viviparity enrichment factor for the results at *K* = 2 as:

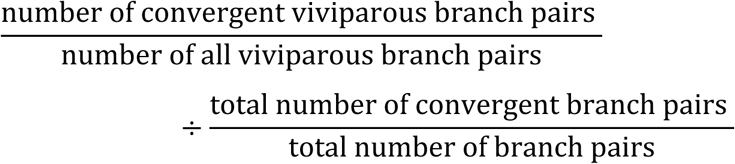

An enrichment factor > 1 indicates a higher proportion of branch pairs show convergent amino-acid substitutions in the viviparous lineages. A contingency table test was then applied for each transcript alignment to test if this difference in proportion is significant. The isoform with the lowest *p* for the enrichment factor was chosen to represent each gene. Similarly, genes with significant enrichment factor > 1 were considered viviparity-enriched CSGs. Empirical FDR analysis was not possible for CSUBST due to computational resource limit. Instead, we applied direct FDR correction on the raw *p* values using the Benjamini-Hochberg method, which is more conservative than empirical FDR.

### Analysis of phylogenetic distance and the frequency of molecular convergence

To test if the number of PSGs shared between two independent viviparous clades are related with their phylogenetic distance, we conducted a corelation analysis. The number of genes shared between two lineages were normalised using Jaccard similarity to account for difference in number of PSGs in each lineage:

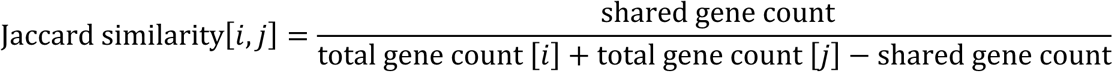

Pairwise phylogenetic distance was calculated based on the species tree using “ape” package (90) in R v4.2.1. Pearson’s corelation analysis was conducted using “cor.test” function in R.

To test if the chance of adaptive convergent amino-acid substitutions for a gene was also related with distance between the branch pairs, we calculated the mean distance of all convergent branch pairs in foreground lineages for each gene. The distance for branch pairs were extracted from the CSUBST result file. Because the distance was estimated based on each gene alignment, we normalised the distance by the mean distance of all nonsignificant branch pairs in the viviparous foreground. Consequently, the null hypotheses (no relationship between the chance of convergence and pairwise phylogenetic distance) would be a normalised branch distance of 1. We calculated the mean convergent branch-pair distance for each gene, and then conducted a Wilcoxon signed-rank test to test if the mean distance is significantly different from 1.

### Functional enrichment analysis

To test if the viviparity-enriched gene sets obtained in this study are enriched in viviparity related biological functions, we conducted GO term enrichment analysis (91) for target gene sets relevant to all protein-coding genes in the reference genome (*E. macularius*). All the protein sequences from the reference genome (GCF_028583425.1_MPM_Emac_v1.0_protein.faa downloaded from NCBI) was used for annotation by eggNOG-mapper v2.1.12 web server (http://eggnog-mapper.embl.de) (92). The eggNOG 5 database (93) was used, taxon was restricted to vertebrates, and all GO annotations were used (including inferred from electronic annotation). Other parameters were set as default.

Annotation results were then processed by custom R script to extract annotated GO term for each gene based on the results of the longest isoform. GO enrichment for biological processes was conducted using “topGO” R package v2.52.0 (94). The default “weight01” algorithm was applied, which can reduce redundancy by accounting for the GO hierarchy and highlight more specific, independent terms. Only GO terms with more than 5 significant genes in the background gene set and 2 significant genes in the target gene set were considered. Enriched GO terms were obtained using *p* < 0.05.

To explore more subtle and broader selection signal enriched for viviparity instead of limited on the significant gene list, we applied GSEA analysis (95) using all genes and the mapped KEGG pathways. Pathways were again extracted from eggNOG-mapper annotations. Pathways with gene number smaller than 10 or larger than 1,000 were excluded. Genes were ranked using -log_10_ (*p* values) for the ones with the viviparity enrichment score >1 and using log_10_ (*p* values) for the ones with the viviparity enrichment score ≤1, so that the ranking scores have both positive and negative values. KEGG pathways were than tested for positive enrichment for viviparity using “fgsea” R package v1.35.6 (96) using 10,000 permutations.

## Acknowledgements

We thank the Evolution & Diversity group from SBOHVM, University of Glasgow for valuable discussions about the project. We thank D. Wu and T. Tang for help in visualisation.

## Author contributions

K.R.E. and O.E.G conceived the project. D. H. and J. S. collected the genome dataset. H. X. and D. H. curated the dataset and conducted the analysis. H.X. and K.R.E. discussed and visualized the results. K.R.E. supervised the project. H. X. wrote the original draft. All authors contributed to the discussion, revision, and final approval of the manuscript.

## Data and materials availability

The genomic data used in this work are all from public resources (see **table S1** for detailed entries). TOGA annotation results for all squamate genomes, summary result files for selection analysis, and custom python and R scripts used in this study are archived and available on University of Glasgow Enlighten [permanent doi with acceptance].

