## Supplementary Material for "Convergent molecular evolution of viviparity across squamate reptiles"

**This file includes:**

Supplementary Figures S1-S8

Supplementary Table legends S1-S15

### Supplementary figures

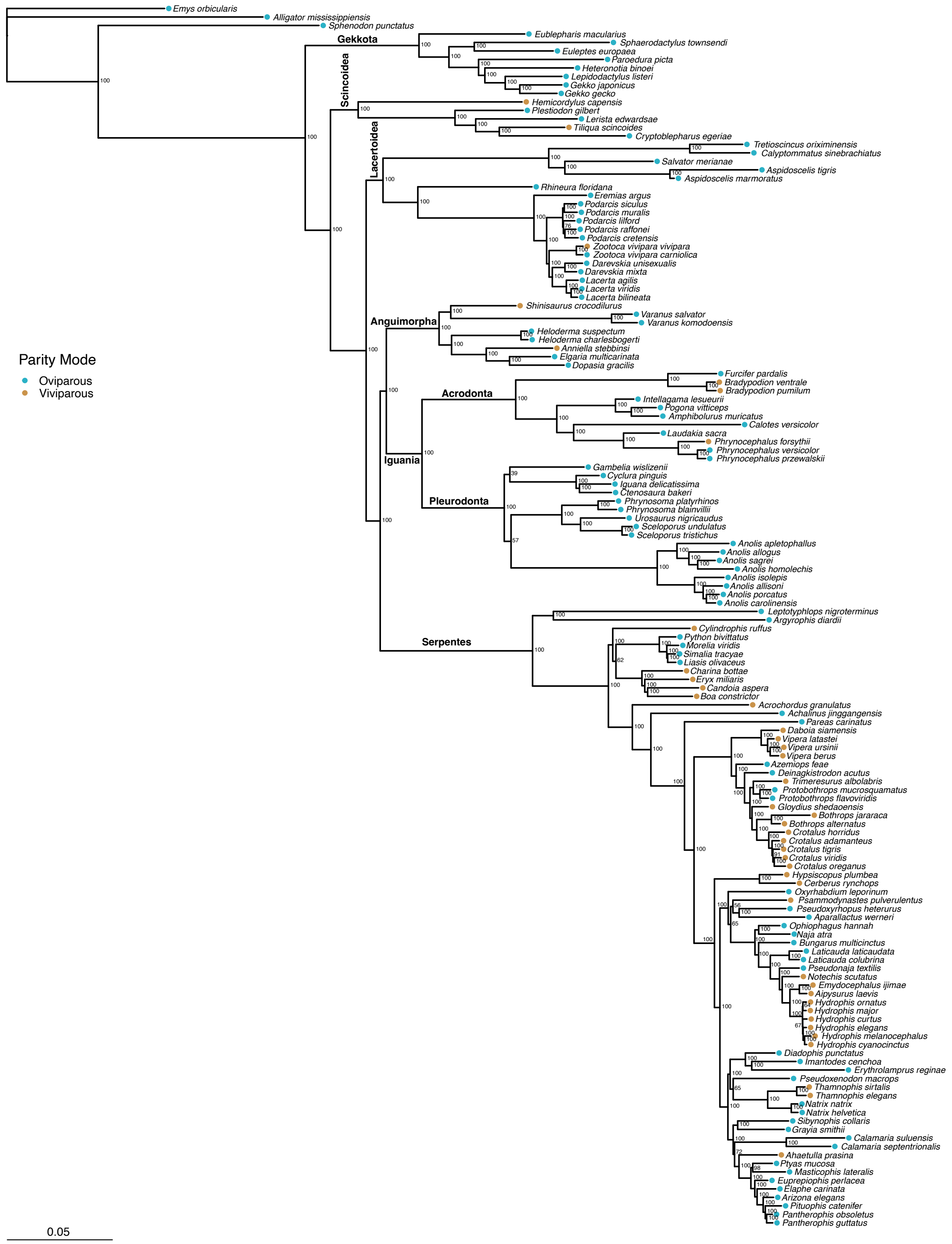


**Figure S1.** Species phylogenetic tree generated in this study. Tree was built using maximum likelihood method based on protein alignment of 1,605 universal one-to-one genes. Numbers on the node represents bootstrap values.

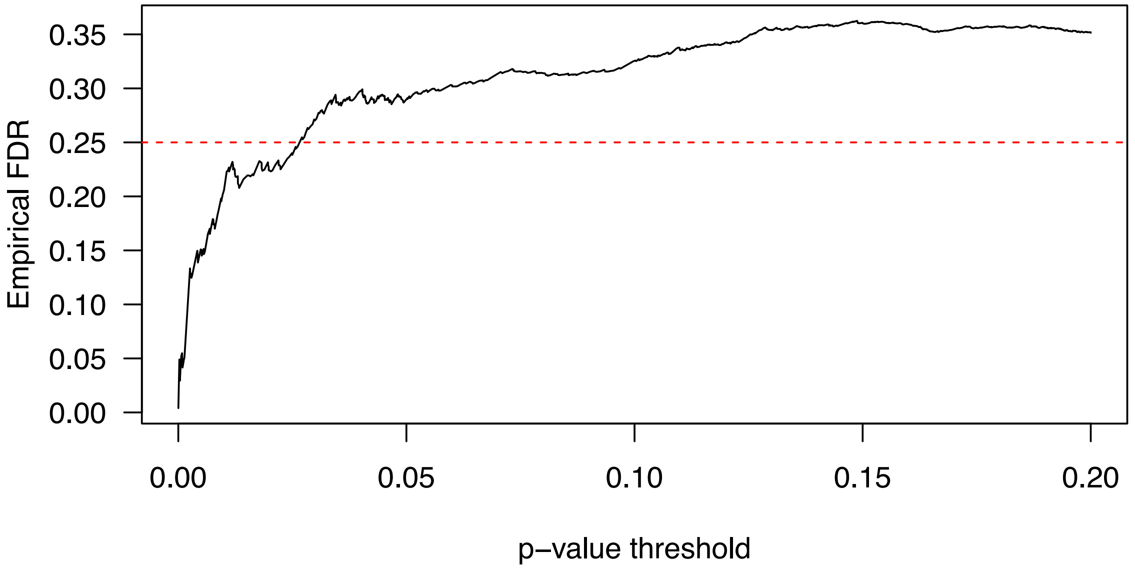


**Figure S2.** The empirical false discovery rate (FDR) obtained from 1,000 permutations for the observed *p* values ≤ 0.2 for the enrichment of positive selection in viviparous lineages. An FDR of 0.25 (red dashed line) was used in this study.

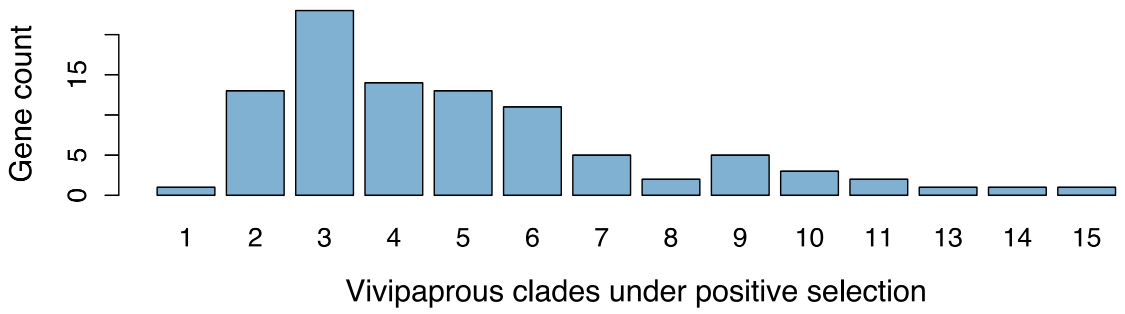


**Figure S3.** Distribution of the number of significant viviparous clades under positive selection for the subset of 95 viviparity-enriched PSGs.

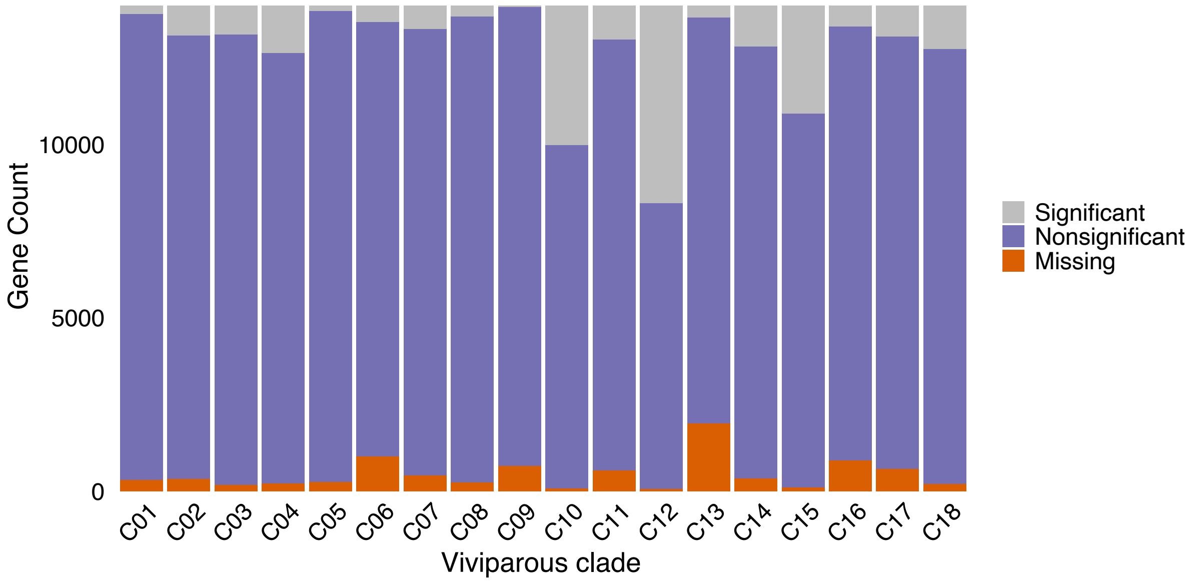


**Figure S4.** The number of genes showing positive selection signal in each viviparous clade for the background gene set (all genes except the 95 viviparity-enriched PSGs). For each clade, the number of genes having "significant" positive selection signal, "nonsignificant" positive selection signal, or is "missing" in the alignment (due to assembly gap or quality control) are shown in stacked bars.

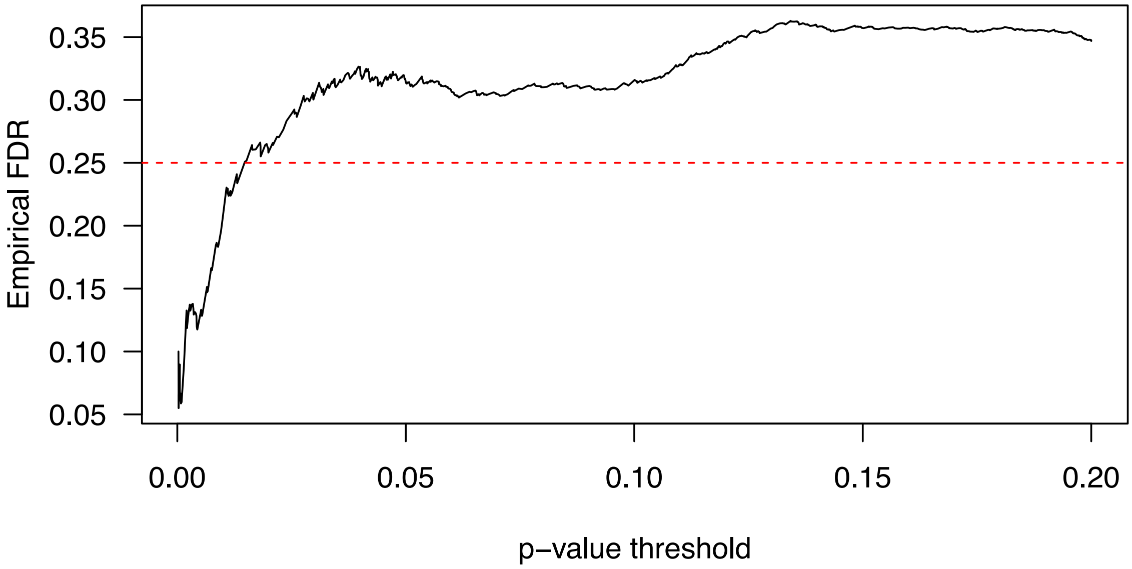


**Figure S5.** The empirical false discovery rate (FDR) obtained from 1,000 permutations for the observed *p* values ≤ 0.2 for the enrichment of intensified selection in viviparous lineages. An FDR of 0.25 (red dashed line) was used in this study.

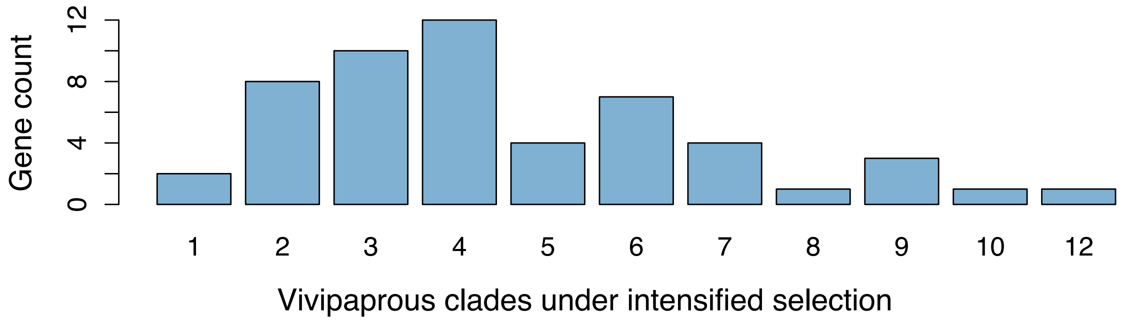


**Figure S6.** Distribution of the number of significant viviparous clades under intensified selection for the subset of 53 viviparity-enriched ISGs.

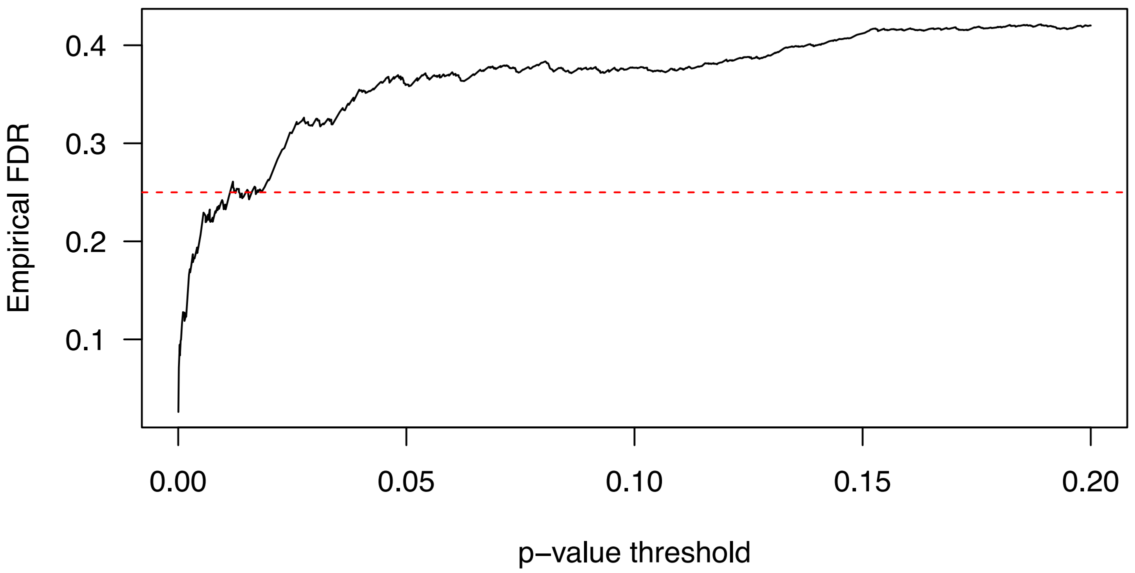


**Figure S7.** The empirical false discovery rate (FDR) obtained from 1,000 permutations for the observed *p* values ≤ 0.2 for the enrichment of relaxed selection in viviparous lineages. An FDR of 0.25 (red dashed line) was used in this study.

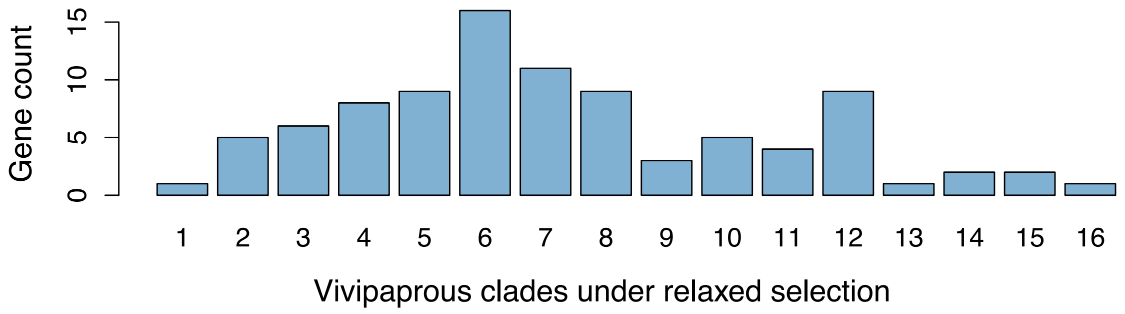


**Figure S8.** Distribution of the number of significant viviparous clades under relaxed selection for the subset of 92 viviparity-enriched RSGs

#### Supplementary tables

**Table S1.** Collected and compiled squamate genome resources by Feb 2024. This list also include an outgroup species from Crocodilia, Testudines, and Rhyncocephalia, respectively.

**Table S2.** Curated and annotated genomes (N = 144) used in this study. BUSCO scores include complete genes (C), complete single gene (S), complete duplicated gene (D), fragmented gene (F), and missing gene (M). TOGA annotations include (in the order of reference-to-query) one-to-one gene (1to1), one-to-many gene (1toM), many-to-one gene (Mto1), many-to-many gene (MtoM), and missing gene (1to0).

**Table S3.** Summary statistics of the 232 genes that show enriched positive selection signal for viviparity. P-values were obtained for a contingency table test for the enrichment factor using Fisher's exact test. Genes are listed in the order of increasing *p* value. FDR thresholds was obtained by conducting 1,000 empirical permutations.

**Table S4.** Summary statistics of the 205 genes that show enriched intensified selection signal for viviparity. P-values were obtained for a contingency table test for the enrichment factor using Fisher's exact test. Genes are listed in the order of increasing *p* value. FDR thresholds was obtained by conducting 1,000 empirical permutations.

**Table S5.** Summary statistics of the 212 genes that show enriched relaxed selection signal for viviparity. P-values were obtained for a contingency table test for the enrichment factor using Fisher's exact test. Genes are listed in the order of increasing *p* value. FDR thresholds was obtained by conducting 1,000 empirical permutations.

**Table S6.** Summary statistics of the 211 genes that show enriched convergent substitutions for viviparity. P-values were obtained for a contingency table test for the enrichment factor using Fisher's exact test. Genes are listed in the order of increasing *p* value. Adjusted *p* was obtained using the Benjamini-Hochberg method.

**Table S7.** Significant GO terms for the 95 viviparity-enriched PSGs. Only terms with at least five genes in the background and two genes in the target gene set were retained.

**Table S8.** Significant GO terms for the 53 viviparity-enriched ISGs. Only terms with at least five genes in the background and two genes in the target gene set were retained.

**Table S9.** Significant GO terms for the 92 viviparity-enriched RSGs. Only terms with at least five genes in the background and two genes in the target gene set were retained.

**Table S10.** Significant GO terms for the 1118 viviparity-enriched CSGs. Only terms with at least five genes in the background and two genes in the target gene set were retained.

**Table S11.** The summary table of whether a gene shows positive selection (1) or no positive selection (0) in each of the 18 viviparous clades (see **Fig. 1** for the clades from C01 to C18). The 14 genes listed are the overlapping genes of viviparity-enriched PSGs and viviparity-enriched CSGs. In other words, these 14 genes show both enriched positive selection and convergent amino acid substitutions in viviparous lineages.

**Table S12**. GESA enrichment results for KEGG pathways under convergent positive selection signal in viviparous lineages.

**Table S13**. GESA enrichment results for KEGG pathways under convergent intensified selection signal in viviparous lineages.

**Table S14**. GESA enrichment results for KEGG pathways under convergent relaxed selection signal in viviparous lineages.

**Table S15**. GESA enrichment results for KEGG pathways under convergent amino acid site substitution signal in viviparous lineages.
